# LightDock goes information-driven

**DOI:** 10.1101/595983

**Authors:** Jorge Roel-Touris, Alexandre M.J.J. Bonvin, Brian Jiménez-García

## Abstract

The use of experimental information has been demonstrated to increase the success rate of computational macromolecular docking. Many methods use information to post-filter the simulation output while others drive the simulation based on experimental restraints, which can become problematic for more complex scenarios such as multiple binding interfaces. We present a novel two-step method for including interface information into protein docking simulations within the LightDock framework. Prior to the simulation, irrelevant regions from the receptor are excluded for sampling (filter of initial swarms) and initial ligand poses are pre-oriented based on ligand input information. We demonstrate the applicability of this approach on the new 55 cases of the Protein-Protein Docking Benchmark 5, using different amounts of information. Even with incomplete information, a significant improvement in performance is obtained compared to blind ab initio docking.

The software is supported and freely available from https://github.com/brianjimenez/lightdock and analysis data from https://github.com/brianjimenez/lightdock_bm5.

## 1 Introduction

Computational tools are essential to predict and describe threedimensional (3D) interactions between biomolecules. In particular, integrative approaches, i.e. data- or information-driven, are broadly used in order to combine experimental data with docking simulations (De Vries et al., 2010, 2015; Russel et al., 2012; Jiménez-García et al., 2013; Rodrigues and Bonvin, 2014; Quignot et al., 2018). In the context of molecular docking, there are still two main challenges: (1) searching the conformational space, especially in the case of highly flexible molecules, and (2) evaluating and selecting near-native poses out of the generated conformers, which is usually referred to as scoring.

LightDock (Jiménez-García et al., 2018) is a multiscale flexible framework for the 3D determination of binary protein complexes based on the Glowworm Swarm Optimization (GSO) (Krishnanand and Ghose, 2009) algorithm that systematically optimizes the generated docking poses towards those energetically more favourable at every simulation step. Here we describe and benchmark an updated implementation of LightDock that now supports the use of information to drive or bias the docking simulation by filtering out swarms, pre-orienting ligand poses based on the available information and biasing the scoring energy upon satisfied restraints.

The results on the benchmark demonstrate a high performance of LightDock when used in combination with additional information. We also explore different scenarios with less accurate or limited information to show the versatility of our approach.

## 2 Methods

Introducing restraints or biases in docking is a powerful mechanism to drive the simulation towards poses that satisfy those restraints (Dominguez et al, 2003). Due to the nature of the LightDock framework, information about interfacial residues can be applied at different levels depending on the availability of information for the receptor, the ligand or both. On the receptor side, we filter out initial swarms that are not in the proximity of the defined restraints (2.1). On the ligand side, we orient initial poses based on randomly selected receptor-ligand restraint pairs (2.2). Finally, we bias the scoring according to the percentage of satisfied restraints (2.3) at every simulation step.

### 2.1. Swarms selection based on receptor residue restraints

LightDock simulations are organized in swarms over the receptor surface. Given an initial number of swarms *S* (by default 400) and residue restraints *R* specified by the user, we select the ten closest swarms to each residue in *R* (Euclidean distance). The set of swarms to be simulated is therefore the union of the different swarms selected for each restraint residue, which is a subset of the initial number of swarms *S*.

### 2.2. Glowworms pre-orientation based on ligand residue restraints

Each glowworm in the swarm encodes a given complex pose. The poses evolve in translational (Cartesian), rotational (Quaternions) and conformational space through an Anisotropic Network Model (ANM) space. The ANM model considers (by default) the ten first non-trivial normal modes calculated on the Ca and further extended to the rest of the atoms. These are included in each glowworm optimization vector to model backbone flexibility of both receptor and ligand molecules.

For each swarm, we select from the set of input restraints, the 10 closest receptor residues with respect to the geometric centre of the swarm (R_c_). Then, we create random receptor-ligand restraint pairs {r, 1} where r ∈ R_c_ and l is a defined restraint residue of the ligand molecule. Finally, we orient each ligand pose using the vector facing the direction given by {r, 1}. **Figure 1** shows the preferred orientation of yellow arrows pointing towards the receptor restraint residues.

**Fig. 1.**
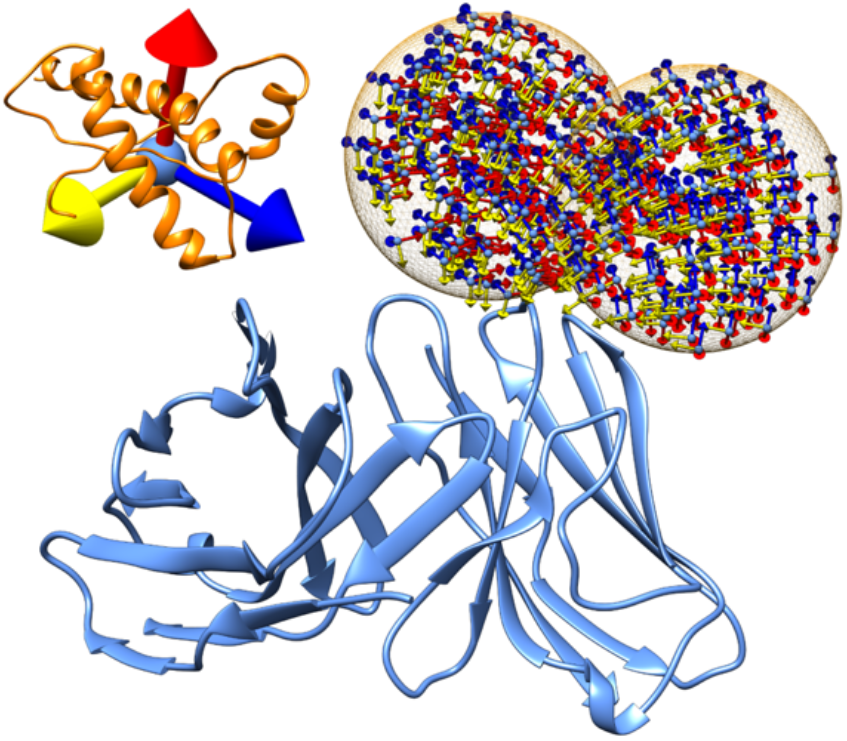
Representation of the filtered and pre-oriented swarms. Representation of two swarms (orange mesh) over the surface of a receptor protein (blue). The initial orientations of the ligands in the swarm are represented using an orthogonal axis.

### 2.3. Score bias according to percentage of satisfied residue restraints

LightDock is somehow agnostic of the scoring function as previously discussed in (Jiménez-García et al., 2018). The overall quality of the simulation will, of course, heavily depend on the capabilities of the selected scoring function to successfully describe the protein docking energetic landscape. In this new implementation, we calculate the intersection between the set of input restraints provided by the user and the set of those in contact for a given pose (3.9Å distance cutoff). The final score ε_f_ (Eq. 1) of the complex is increased by the percentage of satisfied restraints (no penalties if none of the restraints is satisfied).

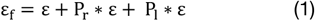

ε is the energy as calculated by the scoring function, and P_r_ and P_l_ are the percentage of satisfied restrained residues of the receptor and ligand, respectively.

## 3 Results

The latest release of LightDock (0.6.0) (Jiménez-García et al., 2019), which now supports the use of information to drive the docking in the format of residue restraints, was tested on the 55 new entries of the Protein Docking Benchmark version 5 (Vreven et al., 2015). We defined various scenarios to demonstrate its versatility as follows:

(1) TI: True interface, defined as those residues at 3.9Å distance from the partner molecule. This is an ideal case where a fully accurate definition of interface residues is available, but no specific contacts are defined.
(2) TI_50_: Half of the residues defined as TI are randomly discarded. This represents a scenario where the amount of information is more restricted but still accurate. On average, 8 residue restraints are selected from the receptor and ligand interfaces.
(3) TI_25_: One fourth of the TI is kept as restraints. On average, this represents 4 residues from receptor and ligand interfaces.
(4) TI_REC_: Only the TI from the receptor is considered as restraints.
(5) PAIR: Only one receptor-ligand residue pair, making a real contact, is used as residue restraints. This case may exemplify a double mutant experiment.

**Figure 2** shows the results for the five scenarios described above together with ab initio docking, which is included as a baseline for comparison. The scoring function used in these LightDock simulations is DFIRE (Zhou and Zhou, 2009). The predictive performance of LightDock when full interface (TI) or half of the residues (TI50) is used remains uniform with 67% and 64% for the Top1 up to 98% in the Top100 for TI and TI50. Interestingly, in the scenario where only one fourth of the interface is used (TI25), the protocol is very robust, with a Top5 performance comparable with the other scenarios (success rate ~80%). Remarkably, for 69% of the cases when only one residue pair is defined (PAIR), LightDock predicts and scores a near-native solution in the Top1. In the scenario where only the contribution of the receptor is taken into account (TI_REC_), while the Top5 performance drops, still a substantial success rate of 67.3% is obtained for the Top100. This last scenario is specially interesting as it directly applies for example to anti-body-antigen docking where antibody information is known beforehand (the CDR loops) but not the epitope on the antigen.

**Fig. 2.**
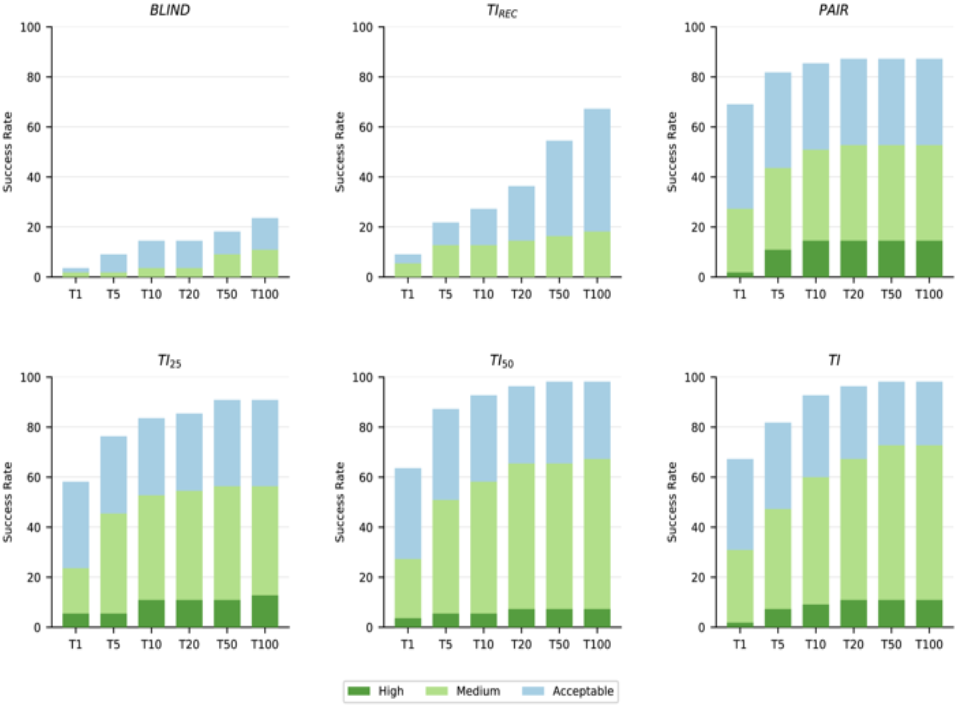
Performance for different scenarios. *Ab initio* docking (BLIND), using only receptor contribution to true interface (TI_REC_),a single residue pair from the true interface (PAIR), using 25% randomly selected residues from the true interface as restraints (TI25), 50% randomly selected interface residues (TI_50_) and using all the residues from the true interface (TI). True interface residues are calculated at a cutoff distance of 3.9Å. Results are presented according to the CAPRI quality criteria (Lensink and Wodak, 2010) and the success rate is defined as the percentage of cases with at least one non-incorrect model within a given Top *N* (*N*= 1, 5, 10, 20, 50 100).

For comparison, results for the full ab initio mode (BLIND) of LightDock are also reported, with top 10 and 100 performance of 14.5% and 23.6%, respectively. These are clearly much lower than any of the other scenario tested here.

## 4 Conclusion

The new version of LightDock offers a powerful tool for modelling protein-protein complexes with high accuracy when some information about interfaces is available. Next to enabling the incorporation of data from mutagenesis and/or bioinformatics predictions, for example, this strategy might also be convenient in scenarios such as limiting the sampling to the solvent accessible loops of a transmembrane protein, or the CDR loops of an antibody. While other FFT-based methods do support a-posteriori filtering, the prefiltering of swarms in LightDock does lead to a reduction of the computation time, which could be used to ensure a denser sampling around the binding region.

